# Phylogenomics of SAR116 clade reveals two subclades with different evolutionary trajectories and important role in the ocean sulfur cycle

**DOI:** 10.1101/2021.05.06.443042

**Authors:** Juan J. Roda-Garcia, Jose M. Haro-Moreno, Lukas A. Huschet, Francisco Rodriguez-Valera, Mario López-Pérez

## Abstract

The SAR116 clade within the class Alphaproteobacteria represents one of the most abundant groups of heterotrophic bacteria inhabiting the surface of the ocean. The small number of cultured representatives of SAR116 (only two to date) is a major bottleneck that has prevented an in-depth study at the genomic level to understand the relationship between genome diversity and its role in the marine environment. In this study, we use all publicly available genomes to provide a genomic overview of the phylogeny, metabolism and biogeography within the SAR116 clade. This increased genomic diversity revealed has led to the discovery of two subclades of SAR116 that, despite having similar genome size (*ca*. 2.4 Mb) and coexist in the same environment, display different properties in their genomic make up. One represents a novel subclade for which no pure cultures have been isolated and is composed mainly of single-amplified genomes (SAGs). Genomes within this subclade showed convergent evolutionary trajectories with more streamlining features, such as low GC content (*ca*. 30%), short intergenic spacers (<22 bp) and strong purifying selection (low d*N*/d*S*). Besides, they were more abundant in metagenomic databases recruiting also at the deep chlorophyll maximum. Less abundant and restricted to the upper photic layers of the global ocean, the other subclade of SAR116, enriched in MAGs, accommodated the only two pure cultures. Genomic analysis suggested that both clades have a significant role in the sulfur cycle with differences in the way in which both clades can metabolize the dimethylsulfoniopropionate (DMSP).

**IMPORTANCE:** SAR116 clade of Alphaproteobacteria is an ubiquitous group of heterotrophic bacteria inhabiting the surface of the ocean, but the information about their ecology and population genomic diversity is scarce due to the difficulty of getting pure culture isolates. The combination of single-cell genomics and metagenomics has become an alternative approach to study this kind of microbes. Our results expand the understanding of the genomic diversity, distribution, and lifestyles within this clade and provide evidence of different evolutionary trajectories in the genome make-up of the two subclades that could serve to understand how evolutionary pressure can drive different adaptations to the same environment. Therefore, the SAR116 clade represents an ideal model organism for the study of the evolutionary streamlining of genomes in microbes that have relatively close relatedness to each other.

## INTRODUCTION

Marine bacterioplankton (phototrophic and heterotrophic) play a central role in the sustainability of marine environments driving main biogeochemical processes as well as primary production at the base of the food chain (1). These microorganisms are believed to be responsible for up to 98% of marine primary productivity (2). In the microbial loop, heterotrophic bacteria are responsible for the assimilation and metabolization of labile dissolved organic matter (DOM) released by photoautotrophs in the aquatic environment (3–5). Variations in the availability and type of nutrients in the pelagic habitat have led to the emergence of distinct trophic strategies. Some models for heterotrophic marine bacteria such as *Alteromonas* (6, 7), *Vibrio* (8) or *Roseobacter* (9) are copiotrophs i.e. grow with high concentrations of nutrients, therefore they are easy to handle in the laboratory and their cells are relatively large. However, these microbes are minorities in the open ocean and play a less remarkable role in the ecosystem. In offshore oligotrophic pelagic habitats, only the transient nutrients discharged from particulate organic matter e.g. in algal blooms or animal eject, provide opportunities for their swift growth (10).

However, the molecular approach first and metagenomics later have proven that the surface ocean microbiome is mostly dominated by prokaryotes (that carry a large part of the load of the planet ecology) belonging to a different lifestyle strategy (oligotrophs) (11–15). Despite their abundance and importance, the bottleneck of getting pure cultures by classical culture-based approaches have slowed down considerably their study. Thus, most of our present knowledge about these largely unknown but essential components of the biosphere and the ocean microbial ecosystem has been derived from metagenomics and single-cell genomics approaches (14, 16–19). Although this kind of microorganisms are extremely diverse, they tend to have a characteristic in common. They have small cells (e. g. *Candidatus* Pelagibacter ubique has 0.12-0.20 µm diameter and a cell volume of only 0.01 µm^3^ (20)), so small that their discovery took longer than most other microbes. The small cell size can be considered a direct effect of the scarcity of nutrients in their environment (21, 22). Most of the ocean water column, contrastingly to soil, sediments or animal bodies, is oligotrophic, i.e. contain highly diluted organic and inorganic nutrients. The microbes that thrive there are mostly oligotrophs that utilize nutrients in very low concentrations. For that, they need to keep a low surface to volume ratio, which translates into very small cells (23). As a consequence, they have highly compacted genomes characterized by i) significant reduction in genome size with highly conserved core genomes and few pseudogenes, ii) short intergenic spacers, iii) low numbers of paralogs and iv) low GC content. These genomic features described as an evolutionary adaptation for more efficient use of nutrients in oligotrophic environments removing non-essential genes were named as “streamlining theory” (23).

Although underrepresented in comparison to these streamlined dominant groups such as the alphaproteobacterial SAR11 clade and the Cyanobacteria *Prochlorococcus* (23), there are many other cosmopolitan lineages of heterotrophic marine bacterioplankton in the global oceans, including SAR116 and SAR86 clade genomes within Proteobacteria, or the Acidimicrobiales within the Actinobacteria (24, 25). Despite playing a central role in the function of marine ecosystems they have received much less attention largely due to the fact that only a few isolates have been isolated or characterized (26). These microbes do not fall into the streamlined category and most of our knowledge about their ecological and genomic role comes from either metagenome-assembled genomes (MAGs) or single-cell genomes (SAGs).

Here, for the first time, we applied an ecogenomic approach to 186 genomes of the SAR116 clade (Alphaproteobacteria), a ubiquitous group of heterotrophic bacteria inhabiting the surface of the ocean, to assess their potential role in the marine pelagic habitat (27). Their relative abundance based on 16S rDNA clone libraries varied in the range of 1 to 17% (26). To date, only two representatives of SAR116 have been cultured and their genome sequenced, “*Ca*. Puniceispirillum marinum” IMCC1322 isolated from surface seawater of the East Sea Basin of Korea (28) and HIMB100, collected off the coast of Hawaii in the subtropical Pacific Ocean (29). The predicted metabolic potential of both strains revealed genes of biogeochemical importance such as proteorhodopsins, carotenoid biosynthesis and carbon monoxide dehydrogenase. In addition, IMCC1322 strain plays an important role in the dimethylsulfoniopropionate (DMSP) cycle via the cleavage pathway to generate dimethylsulfide (DMS) in the surface waters of the oligotrophic ocean (30). Although several metagenomic studies of marine samples have obtained MAGs from this group (18, 31), recently their number has increased by *ca*. 100 new genomes coming from a large library of planktonic bacterial and archaeal SAGs collected from tropical and subtropical epipelagic ecosystems. This study has revealed a new perspective on the genomic complexity of the marine microbiome (19). The increased genomic diversity within this group has led to the discovery of two subclades of SAR116, that coexist in the same environment, but appear to be subjected to different evolutionary pressures in their genome make up. The new subclade that emerged from the improved phylogenomic classification showed genomic features similar to streamlined genomes without genome size reduction. Despite genomic differences, metabolic reconstruction revealed a photoheterotrophic lifestyle with several genes involved in the metabolism of inorganic and organic sulfur compounds. We detected genes in the oxidation of sulphite and thiosulphate in both SAR116 subclades. In addition, we found marked differences in the degradation of the organic DMSP; while the isolate genomes and their closest relatives rely on DMSP lyase to degrade it to DMS, the novel subclade encoded exclusively genes involved in the demethylation pathway which produces (methylsulfanyl)propanoate (MMPA). Our data suggests that SAR116 might play a key role in the sulfur cycle in the surface ocean.

## RESULTS AND DISCUSSION

### Phylogenomic characterization of the SAR116 clade

A total of 186 genomes were downloaded from publicly available databases putatively classified as members of the SAR116 clade (Based on NCBI classification accessed in August 2020; see Material and Methods), which includes only two cultured representatives (IMCC1322 and HIMB100) together with 120 SAGs and 64 metagenome-assembled genomes (MAGs) that met the established quality criteria of ≥ 50% completeness and ≥ 5% contamination i.e medium to high-quality draft genomes (32) (Table S1). Phylogenomic analysis using a concatenation of 258 single copy marker proteins showed that SAR116 genomes clustered into two major subclades with four different families (two per subclade) (Fig 1A). Based on GTDB classification (33), these four families were placed within the Puniceispirillales order (Table S1). The two pure culture representatives were placed in the same family (Puniceispirillaceae) that together with family UBA1172 clustered within one of the subclades characterized by containing a higher proportion of MAGs (60 MAGs and 43 SAGs) (Fig 1A and 1B). On the other hand, the other subclade, composed of families AAA536-G10 and GCA-002684696, was represented mostly by SAGs (#85) including only 4 MAGs (Fig 1A and 1B). Most of these SAGs (79 of the 85 genomes) come from a large collection of genomes released from the surface (epipelagic) ocean in tropical and subtropical latitudes (19) (Table S1). Therefore, this intrinsic difficulty to obtain pure cultures and to reconstruct genomes from metagenomes of this new subclade has meant that its genomic diversity has been hidden until now with the advance in single-cell genomics. Clustering based on pairwise average nucleotide identity (ANI) (Fig S1) revealed groups of genomes within each family with ANI values of *ca*. 70%, which placed these strains likely as different genera, named A to D for simplicity (Fig 1A). In the end, we were able to distinguish two subclades, four families and ten putative genera within the SAR116 clade (Fig 1A and Table S1).

**Fig. 1:**
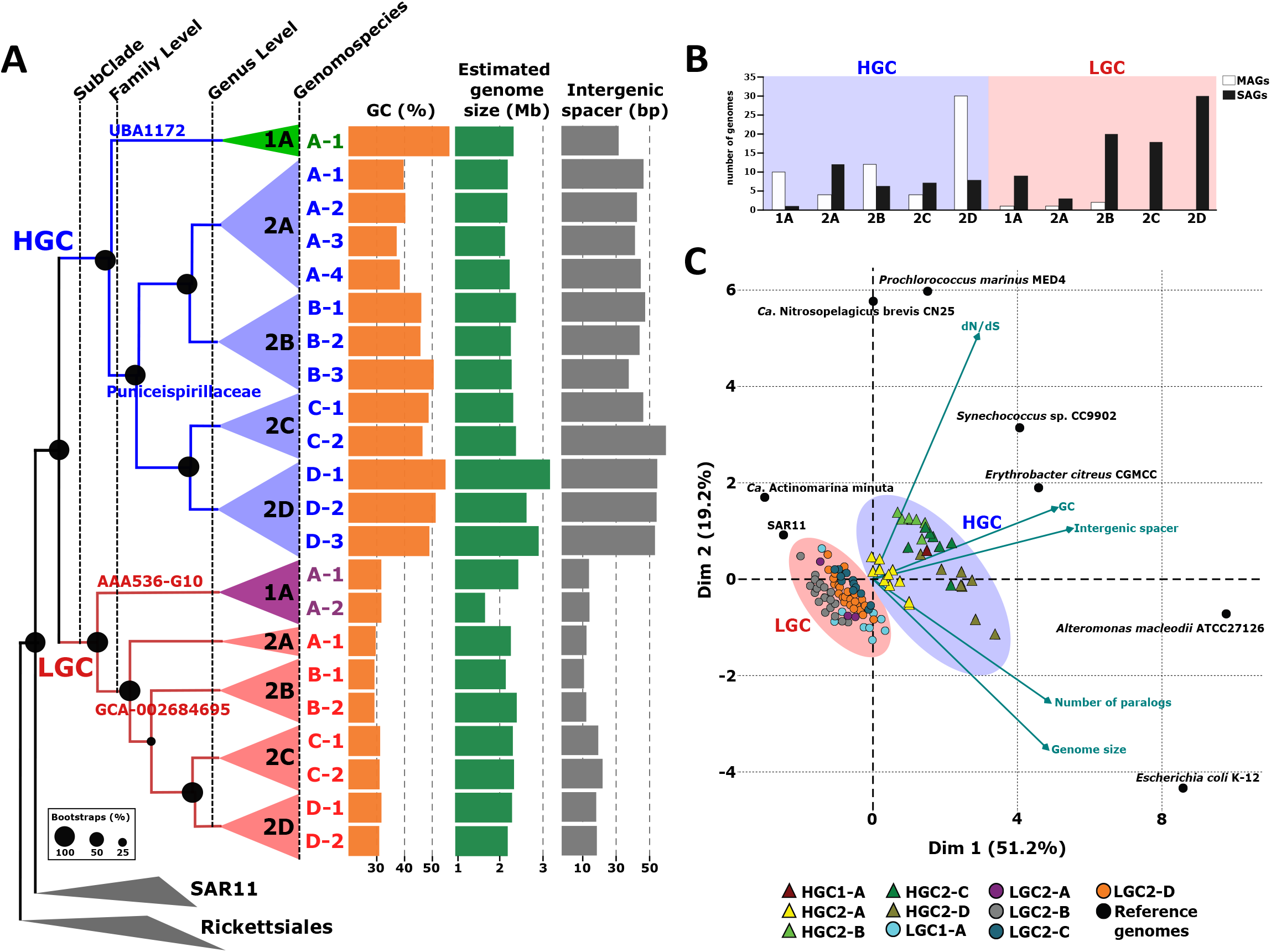
(A) Phylogenomic analysis of all SAR116 genomes available using a total of 258 concatenated conserved proteins to generate a maximum likelihood tree. The branches have been colored according to the subclade to which they belong [Blue, High GC (HGC) and Red, Low GC (LGC)]. The genomes of nearby orders SAR11 and Rickettsiales were used as outgroup. GC content, together with estimated genome size and intergenic spacer are plotted next to the tree. (B) Number of SAGs and MAGs belonging to each genospecies within the HGC and LGC subclades. (C)

### Differential genomic features of the SAR116 subclades

Once the phylogenomic classification of the whole clade was established, genomic features were evaluated for each group. Since most of them were incomplete genomes, we calculated the estimated genome size, the GC content (%GC) as well as the intergenic spacer length (Fig 1A and Table S1). Interestingly, we found a significant variation of the GC content between the two subclades. While the subclade containing pure culture representatives (Puniceispirillaceae and UBA1172 families) showed a wide range of values from 38 to 55%, (average *ca*. 47%), this value was consistent across all genera in the new subclade (*ca*. 30%) (Fig 1A and Table S1). Based on these differences, we tentatively named the two subclades as High GC (HGC) and Low GC (LGC) (Fig 1A). The lower GC content has been suggested to be a natural adaptation in nitrogen-limited environments such as open ocean regions (23). In fact, we observed changes in the amino acid usage between both groups. LGC subclade showed higher prevalence for basic amino acids such as Asparagine and, Lysine with only one N atom in side-chains. However, members of the HGC group had a higher frequency of Arginine (3 N in side-chain) (Table S2). In addition to the GC content, we observe a significant variation in the intergenic spacer length. While in HGC they were between 35 and 59 bp (median 49), in none of the genera of the LGC subclade median spacers were longer than 22 bp, with values as low as 11 bp in the case of LGC2B (Table S1 and Fig 1A). Remarkably, the estimated genome size of the genomes was similar in all genera from both subclades (*ca*. 2.4 Mb), with the only exception of the genus HGC2D, which showed a genome size higher than the rest with an average of 3.2 Mb (Table S1 and Fig 1A). Likewise, this genus also exhibited high values for both GC content and intergenic spacer sizes. As a consequence of the smaller size of the intergenic space, genomes within the LGC have on average more than 100 genes per Mb of genome (Table S3).

These genomic features suggested that members within the LGC subclade are undergoing a streamlining process without genome reduction. For that reason, we studied other characteristic genomic parameters that have been proposed to be relevant in the streamlined genomes such as selective pressure and the number of paralogs (34–37). Microevolution was measured as the ratio of nonsynonymous to synonymous polymorphisms (d*N*/d*S* ratio). We found that the median d*N*/d*S* value was *ca*. 0.09 for LGC, this value was comparable to the better known marine SAR11 clade (34) and suggests a strong purifying selection acting on the genome evolution of this subclade (Table S3). Within the HGC subclass, we observed much more variable values. While the genus HGC2A showed similar values to the LGC (0.08), in the other genera within the HGC we found markedly higher median d*N*/d*S* (*ca* 0.15) (Table S3). However, the number of paralogs (*ca*. 170) was consistent across genera in both subclades (Table S3).

In order to put these genomic features into perspective, we compared these groups with a collection of reference marine microbes with different ecological strategies (Table S3 and Fig 1C). Despite the divergence, genomes within the LGC subclade showed consistent genomic parameters, some of them (GC content and d*N*/d*S* ratio) typical of well-studied streamlined genomes such as SAR11 or *Ca*. Actinomarina minuta (35) (Table S3 and Fig 1C). The median intergenic distance was higher than these two microbes, although it was slightly lower than other marine microbes with streamlined genomes such as marine ammonia-oxidizing thaumarchaeon “*Ca* Nitrosopelagicus brevis” and the cyanobacterial *Prochlorococcus marinus* CCMP1986 (Table S3 and Fig 1C). However, the estimated genome size was double that of all these reference genomes. On the other hand, the HGC group shows multiple genomic evolutionary trajectories with features more similar to the marine copiotrophic heterotrophs such as *Erythrobacter* and *Alteromonas* or the cyanobacterium *Synechococcus* sp. CC9902. The case of the HGC2A group is outstanding in displaying an intermediate trend with strong purifying selection and lower GC more similar to LGC (Table S3 and Fig 1C). In addition, like the LGC groups, HGC2A had a higher proportion of genomes recovered by single-cell genomics (Fig 1B).

### Ecological distribution (metagenomic recruitment)

The differential genomic features observed between both subclades could be related to adaptations to specific ecological niches. Therefore, we analyzed the distribution patterns using metagenomic read recruitment analysis in the large global dataset from the *Tara* Oceans Project (16).

First, we analyzed the relative abundance of all the genomes against their occurrence in the metagenomics samples which allowed for the determination of several genomospecies i.e groups of genomes with close phylogenomic relationship and similar relative abundances within the same geographical locations (35, 38). We were able to differentiate 22 genomospecies (Table S1 and Fig 1A). The minimum pairwise ANI value among these ecogenomic units of classification was *ca*. 85%. The results showed that SAR116 clade microbes were found exclusively associated with the upper layers of the epipelagic zone. None of the genomospecies was present in the cold-water stations of the Southern Ocean or in mesopelagic zones (>200m) (Fig 2A). While HGC members were only found in surface waters, LGCs showed a broader distribution, being present at a higher number of stations and depths, which suggests adaption to a wider range of conditions (Fig 2A). For instance, genomospecies LGC1-A1 and LGC1-A2 recruited in the highest number of stations from surface and deep chlorophyll-maximum (DCM) (Fig 2A). While genomospecies B1, B2 within the HCG2 and A1, A2, B1, C1 and D1 from the LGC could be considered the most cosmopolitan, present in several oceanic provinces from 30°N to 30°S, other genomospecies presented predilection for specific regions such as the Mediterranean Sea (HGC1-A1 and HGC2-A2) and Pacific Ocean South-East (HGC2-A1 and LGC1-A3) (Fig 2A). The highest recruitment values (>20 RPKGs) within the HGC subclade corresponded to the HGC1-A1 and HGC2-D1 genomospecies at the same station in the eastern Mediterranean Sea (TARA_025). Regarding the other subclade, LGC2-C1 presented the highest recruitment values in stations TARA_004 (ANE; Atlantic North East) together with TARA_094 and TARA_096 from temperate waters in the South Pacific Ocean (Fig 2A).

**Fig. 2:**
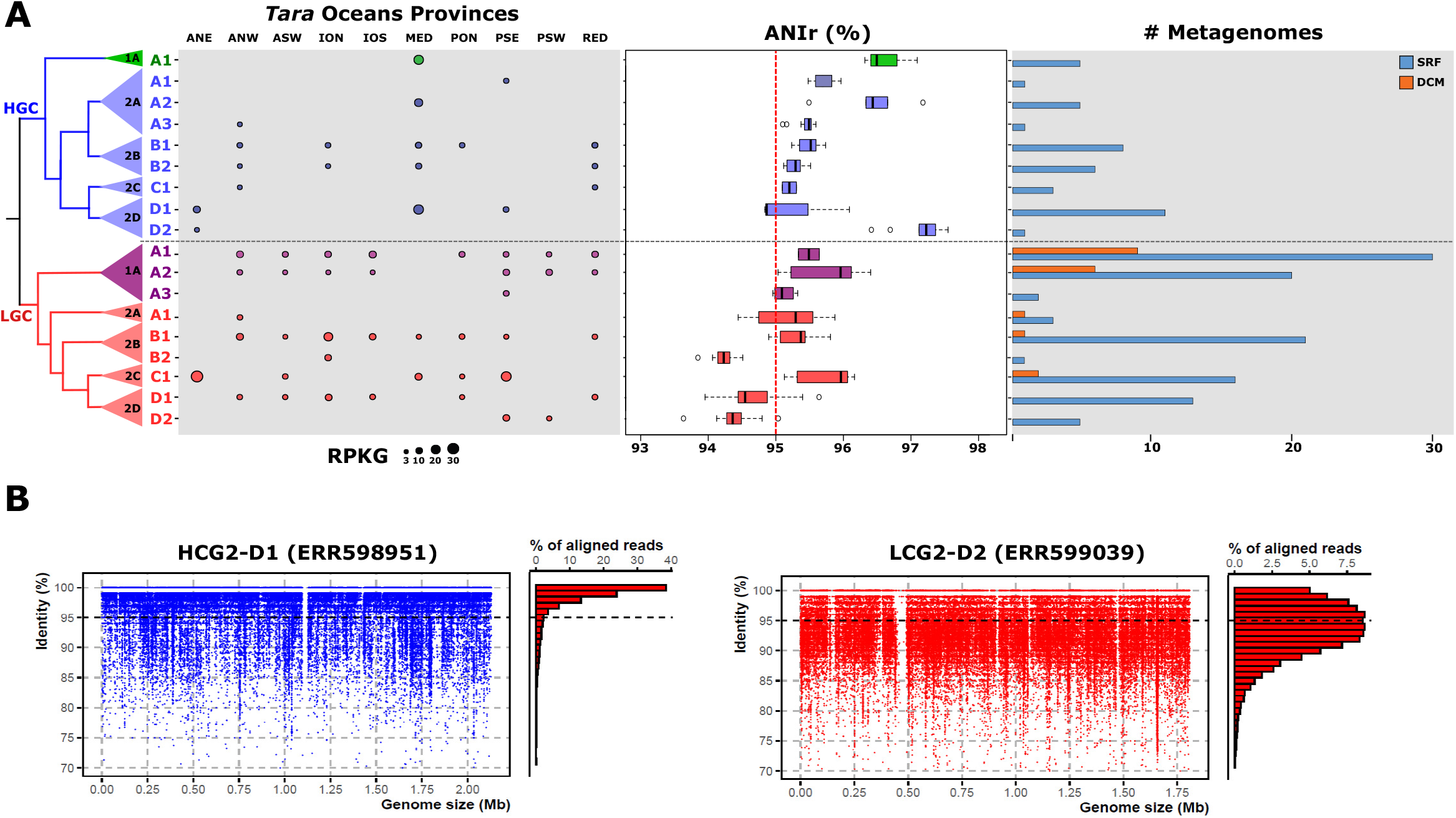
(A) Relative abundance (measured in RPKG) of SAR116 genomospecies in *Tara* Ocean metagenomes. Box plot in the middle indicates the average nucleotide identity based on metagenomic reads (ANIr) among SAR116 genomospecies. Occurrence of SAR116 genomes within *Tara* stations is shown on the right. Bars indicate the number of metagenomic samples where genomes recruit at least three RPKG (presence). A maximum likelihood phylogenomic tree of the SAR116 clade is shown on the left. Box plot and dots from the recruitment were colored according to the different families following the patterns in figure 1. (B) Linear recruitment plot of the representative genomes for HCG2-D1 and LGC2-D2 genera. Each blue dot represents a metagenomic read. The histogram on the right shows the relative percentage of aligned reads in intervals of 1% identity. The black dashed line indicates the species threshold (95%).

In order to examine the intrapopulation sequence diversity, we used the metagenomic recruited reads to determine the read-based average nucleotide identity (ANIr). Most genomospecies in both subclades (HGC and LGC) showed a median ANIr value of *ca*. 95% (species threshold). None of the genomospecies within the HGC presented a lower value but genomospecies HGC1-A1, HGC2-A2 and D2 showed lower intrapopulation sequence diversity (ANIr >96%). These genomospecies could be considered endemic to the Mediterranean Sea and the station TARA_004 (located at the connection between the Mediterranean and the Atlantic Ocean). Therefore, it could suggest a more recent divergence of these groups adapted to the special conditions of the Mediterranean such as limiting P concentration. A similar example has already been described in the SAR11 genomospecies Ia.3/VII, which also showed a preferential presence in the Mediterranean (34). However, three LGC subclade genomospecies (LGC2-B2, LGC2-D1 and D2) showed higher intra-population diversity which could indicate higher ecological persistence over time of these populations (Fig 2A) (39). This is reflected in the linear recruitment plots of these genomospecies (LGC2-D2) with a minimum alignment identity threshold located at *ca*. 85% and HGC2-D1 whose pattern could be associated with a less diverse population (*ca*. 97%) (Fig 2B).

The linear recruitments revealed the presence of metagenomic islands in two genomospecies (LGC1-A1 and LGC2-C1) belonging to different families within the LGC subclade in metagenomic samples from different locations (Fig S2A and S2B). The results showed a highly hypervariable region that was always preserved. The location of the island was conserved among the genomes within the same genomospecies. Detailed analysis of the gene content showed that they are involved in synthesizing the outer glycosidic envelope of the cells (Fig S2C). This high diversity has been explained because they are important phage recognition targets (40).

### General Metabolic features within SAR116 HGC and LGC genomes

The isolation and sequencing one decade ago of two bacterial strains, IMCC1322 and HIMB100 (28, 29), shed light on the physiology and metabolic potential of the SAR116 clade in the oceans. Here, with the increased genomic diversity of SAGs and MAGs, we have expanded the knowledge of this ubiquitous marine group. Given the incomplete nature of SAGs and MAGs, we used the pangenome as a unit to analyse the metabolism against several functional databases (see methods). We included in the comparison the genome of the pure culture IMCC1322, which was phylogenomically classified into the HCG2C (Table S1). Most of the results are in agreement with previous metabolic reports (28, 29) (Fig 3A). Both HGC and LGC subclades are aerobic, chemoorganotrophic microorganisms, encoding enzymes for the three common glycolysis pathways (Embden-Meyerhof-Parnas, Entner-Doudoroff, and pentose phosphate), although as reported from the pure cultures (28, 29), all genomes of both subclades lack 6-phosphofructokinase (*pfk*A); the tricarboxylic acid cycle (TCA cycle); and the complexes I to IV involved in the electron transport chain (ETC). However, in the latter, some differences arose among subgroups. Complex II succinate dehydrogenase could not be detected within the genus LGC2C (18 genomes).

**Fig. 3:**
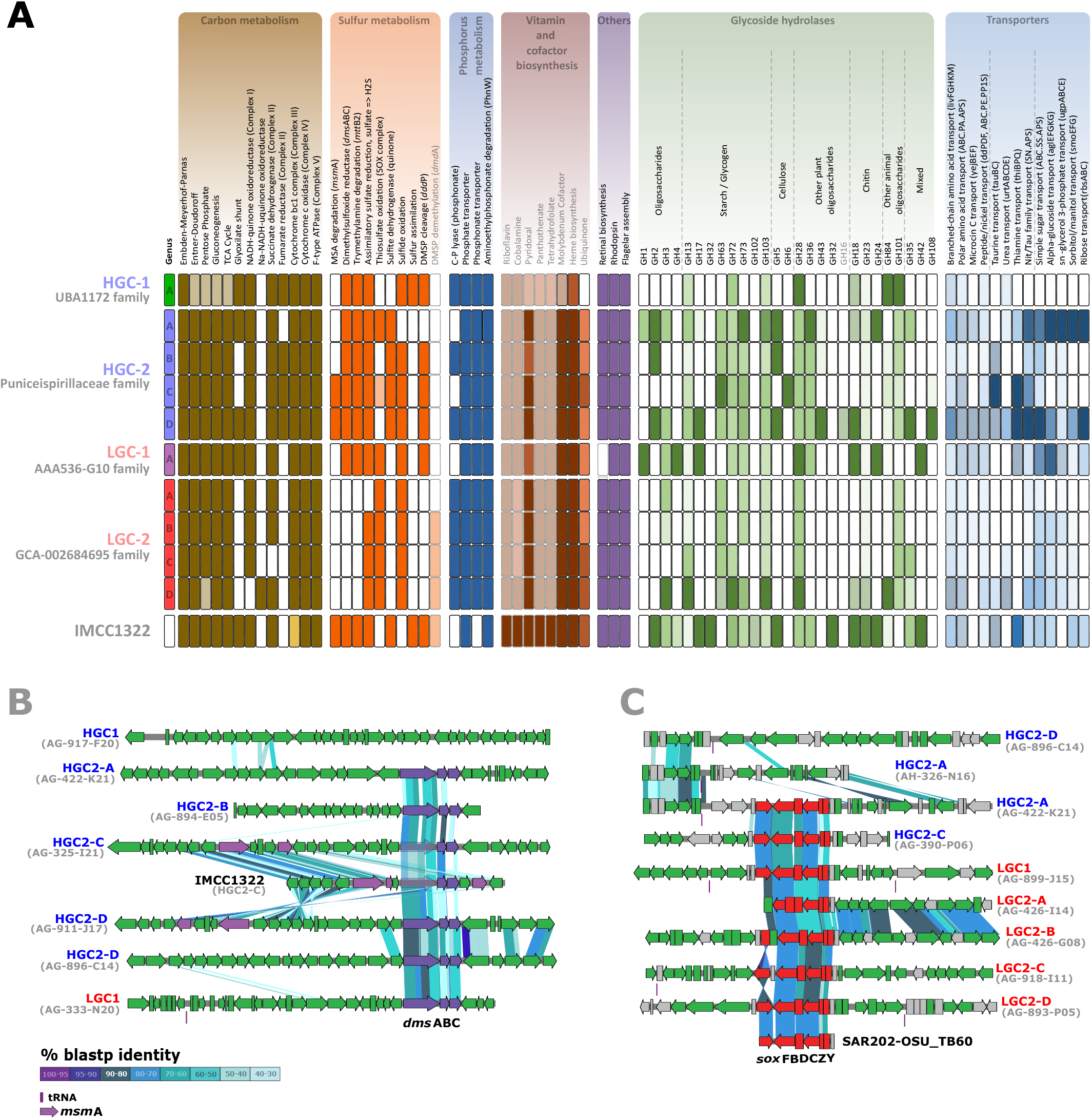
A. Inferred metabolism of the ten SAR116 genera (grouped by family) based on the KEGG database. Ca. Puniceispirillum marinum (IMCC1322) was added for the comparison. B. Genomic alignment (in amino acids) of the *dms*ABC and *msm*A genes found in SAR116 genomes. C. Genomic alignment (in amino acids) of the *sox* operon found in SAR116 genomes. The fragment of SAR202-OSU_TB60 was added for the comparison as the closest relative.

Besides, while the most common version of the complex I detected was the H^+^-<u>N</u>ADH <u>u</u>biquinone <u>o</u>xidoreductase (*nuo*) operon, we detected a horizontal gene transfer (HGT) event within the genomes of LGC2C and LGC2D, on which the *nuo* operon was replaced with the sodium equivalent Na+-pumping NADH:quinone oxidoreductase (*nqr*) operon, being the closest relative to this complex the methylotrophic bacteria HTCC2181 (67.52% average amino acid identity) (Fig S3A). It has already been reported that multiple HGT events have allowed the dispersal of this operon among different bacterial lineages (41). In fact, there is still reminiscence of a gene belonging to the *nuo* cluster (*nuo*L) in these genomes immediately adjacent to the *nqr* operon which is not present in the HTCC2181 genome (Fig S3A). The use of sodium ion transport to generate an electrochemical potential that can be used both for ATP synthesis and also as a primary sodium pump to maintain ionic homeostasis could be an evolutionary advantage in the marine environment. In marine bacteria of the phylum Marinimicrobia the presence of these different versions of respiratory complex I have been correlated with improved ecological adaptation to discrete niches (epipelagic and mesopelagic environments) (42).

The glyoxylate shunt (GS), a two-step metabolic pathway that serves as an alternative to the TCA cycle was only detected in some genera of the HGC subgroup (HGC1, HGC2A, HGC2D) and LGC1. In addition, we detected marked differences in the acquisition and degradation of multiple sugar compounds. Overall, families of glycoside hydrolases (GHs) involved in the degradation of simple and complex oligosaccharides, such as glycogen, cellulose or chitin, and sugar transporters (Fig 3A) were detected in all subgroups, although it is remarkable the elevated number of GH families within genera HGC2 and LGC1. Contrastingly, the low numbers of these degradative enzymes within LGC2 and HGC1 may indicate different ecological strategies degrading organic carbon sources (e.g., cellulase was only detected in HGC2).

Regarding the metabolism of amino acids and vitamins, they all carried the necessary genes for biosynthesis of the twenty common amino acids (data not shown) and the vitamins B2 (riboflavin), B5 (pantothenate) B6 (pyridoxal), B9 (folate), B12 (cobalamin), the molybdenum cofactor, and the heme group (Fig 3A). Remarkably, functional annotation of proteins indicated that instead of using the aspartate 4-decarboxylase, involved in the transformation of aspartate to alanine, they synthesise the latter via the enzyme 2-aminoethylphosphonate aminotransferase (*phn*W) from pyruvate and phosphonate (43, 44).

Lastly, we analysed the presence of some ecologically relevant features. Most of the newly described genera, except LGC1 and HGC2A and B, encoded for genes involved in the acquisition and degradation of phosphonates from seawater. Some regions, such as the Mediterranean or Sargasso seas are depleted in phosphate, organisms inhabiting these places need access to other P-compounds (e.g, phosphonates) to grow and/or survive (38, 45). All of them encoded for the synthesis of a proteorhodopsin. Amino acid sequence analysis indicated that all of them were proton pumps (DTE motif, (46)) and most of them (90 out of 91) absorbed in the blue spectrum. Next to the proteorhodopsin (co-located on the same strand), it is found the gene cluster involved in the synthesis of retinal (Fig S3B). The position of these genes varies between HGC and LGC, and among genera within the high GC groups, which could suggest several independent acquisition events after a common ancestor (Fig S3B). However, in all members of the LGC subclade, the gene coding for isopentenyl diphosphate isomerase (*isp*A) is not present. Remarkably, LGC1 (9 genomes) is the only group that lacks the retinal biosynthesis operon (Fig S3B). This genomic deletion forces the bacterium to retrieve retinal from the environment, like many other marine streamlined organisms (35, 47, 48). Despite the different evolutionary trajectories in terms of genomic architecture, at the functional level, both subclades appear to have many similarities including the absence of essential genes in certain pathways suggesting that they have evolved from a common ancestor.

### Contribution of SAR116 to the sulfur cycle in the ocean

The ocean represents a major reservoir of sulfur (mainly in the form of sulphates) on Earth (49). In this environment, the water column can be considered as a heterogeneous habitat, formed by many kinds of microorganisms that interact with the sulfur cycle. For instance, photosynthetic eukaryotes can reduce sulphate to assimilate it into reduced organic sources. Some bacteria can couple sulphate respiration to degrade organic matter in the absence of oxygen, such as the minimum oxygen zones (50). Conversely, other prokaryotic groups can oxidize inorganic and organic sulfur to produce energy (51).

Functional inference of SAR116 genomes showed that this clade plays a key role in the sulfur cycle (Fig 4). Reduced organic sulfur, in the form of DMSP, an organosulfur compound produced by phytoplankton as compatible solute (52) can be degraded into DMS gas, one of the main sources of sulfur in the atmosphere and reduced sulfur (53, 54) and acrylate by the activity of a DMSP lyase. We found two types of DMSP lyases, *ddd*L and *ddd*P (Fig 4). DMS can be biotically transformed to dimethyl sulfoxide (DMSO) by the enzyme DMS monooxygenase (*dmo*AB), or reduced again to DMS in anaerobic conditions (55) by the enzyme DMSO reductase (*dms*ABC). There is an alternative route to degrade DMSP, which involves the demethylation of DMSP to produce 3-(methylsulfanyl)propanoate (MMPA) by the activity of the enzyme dimethylsulfoniopropionate demethylase (*dmd*A). This is the first step to assimilate sulfur from DMSP into biomass. Some bacteria, such as *Alteromonas macleodii* and *Ruegeria pomeroyi* can continue this pathway to produce acetaldehyde plus methanethiol (*dmd*BCD genes) (56).

**Fig. 4:**
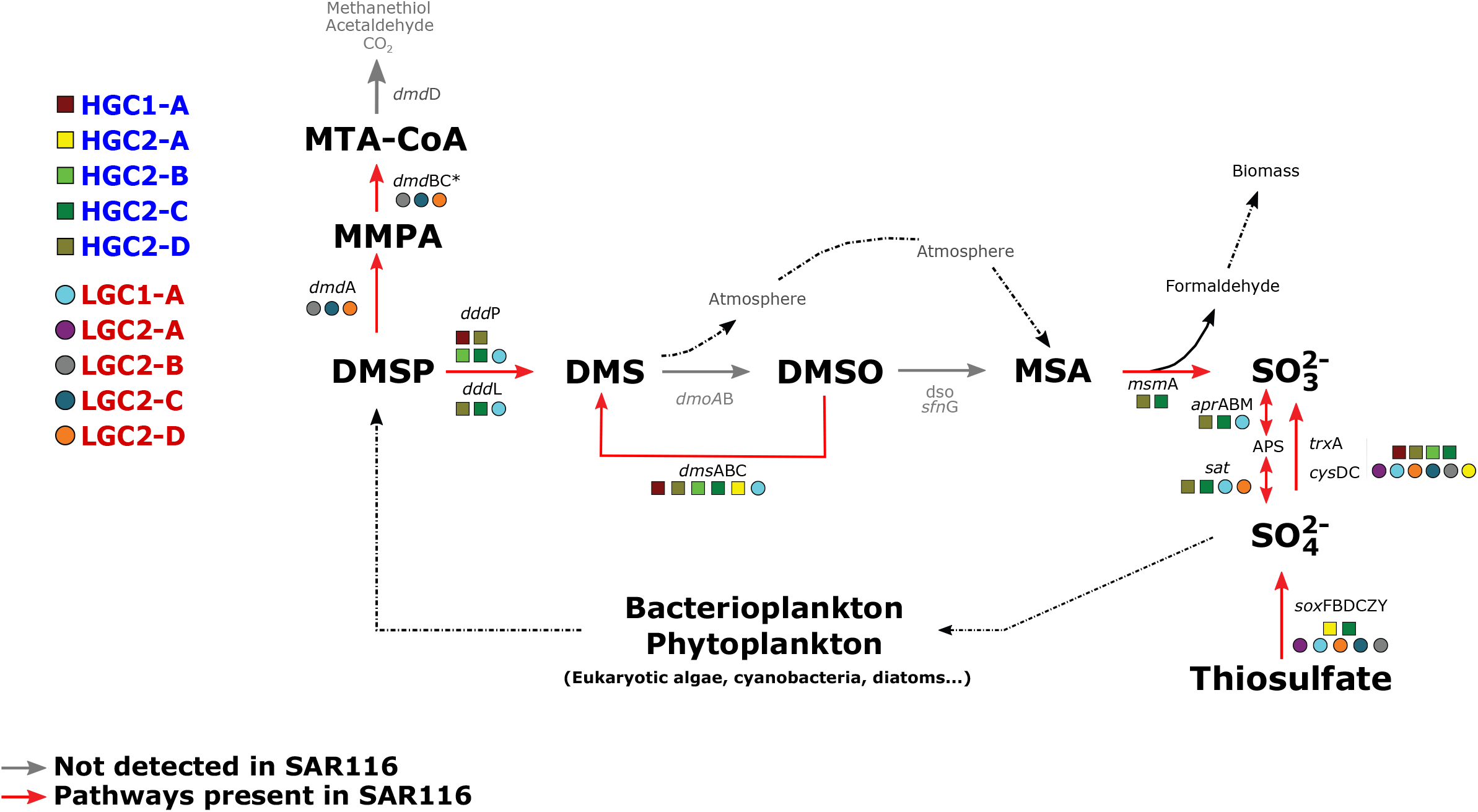
Representative view of the metabolic features found in the different genera of SAR116 related to sulfur cycling. The red lines show the pathways present. Circles and squares indicate genera within the LGC and HGC subclades, respectively.

Figure 4 shows a clear differentiation in the degradation of DMSP by the two SAR116 subclades. Genes involved in the generation of DMS (either through degradation of DMSP by means of DMSP lyases or by reduction from DMSO, *dms*ABC genes) were detected only in the genomes of the HGC2, and LGC1 subgroups (Fig 3B), while the demethylation pathway (*dmd*A) was exclusively detected on LGC subclade (LGC2 genera). Regarding the rest of the genes involved in the degradation of MMPA to methanethiol, we found homologs to *dmd*B and *dmdC* with low identity (*ca*. 40%), but not for *dmd*D. This same pattern has already been described in SAR11 suggesting that the function of this gene (*dmd*D) could be replaced by other non-orthologous isofunctional enzymes (56). It is remarkable that the main pathway to degrade DMSP, found in many epipelagic microorganisms (53) seems to be less relevant in the SAR116 clade. In fact, previous reports already indicated that this clade was the dominant dddP-containing bacteria in the Pacific Ocean (30). DMSO can be further metabolized to methanesulfonate (MSA), which is in turn cleaved to formaldehyde and sulphite by the methanesulfonate monooxygenase (*msm*A). Again, we could detect the MsmA protein in the genomes HGC-2C and 2D, close to the *dms*ABC gene cluster (Fig 3B). In fact, a SAR116 bacteriophage codes for the *msm*A as an auxiliary metabolic gene (57).

Lastly, SAR116 clade codes for several genes involved in sulfur oxidation systems, including the adenosine-5’-phosphosulfate reductase (*apr*ABM) and sulfate adenylyltransferase (*sat*), which catalyse the oxidation of sulphite to sulphate, but only in the genera HGC-2C, HGC-2C and LGC1 [LGC2-D only codes for the *sat* gene (Fig 4)]; as well as the oxidation of thiosulphate by the *sox* operon, widely distributed among LGC1 and LGC2 SAR116 groups, but also detected in HGC2A and HGC2C (Fig 3C and 4). Previous studies demonstrated the presence and activity of sulfur-oxidizing chemolithoautotrophs to use reduced sources of sulfur (e.g. SUP05 and OM252 clades) in anaerobic waters (58, 59), but also in the photic aerobic water column in which *sox* genes are common (18, 60, 61) for energy generation, sometimes coupled to inorganic carbon fixation (62). In this sense, it seems that SAR116, like many other marine prokaryotes (63, 64), may be capable of generating energy from the oxidation of inorganic sulfur on surface waters. Thus, although both subclades appear to be important players in the sulfur cycle in surface ocean waters, the LGC subclade may have an advantage for their capability to demethylate DMSP. The LGC1 group despite its streamlined genome seems to have a higher metabolic versatility than the rest of the LGC group, more similar in this sense to the HGC members, not only in sulfur metabolism but also with a higher richness of both GHs and transporters (Fig 3) which could be one of the reasons for its abundance at DCM (Fig 2).

## CONCLUSIONS

In this study, we have characterized members of the SAR116 clade, an important alphaproteobacterial group of marine heterotrophic bacteria. The enrichment of databases with genomes from single-cell genomics has made it possible to explore in-depth the diversity of this group because of the difficulty in obtaining large numbers of pure cultures using standard methods (to date there are only two pure cultures) and the scarcity and low reliability of MAGs. Phylogenetic analysis suggests that this group of aerobic and chemoorganotrophic microorganisms consists at least of two subclades, four families and ten genera. A new subclass widely represented by SAGs showed genomic characteristics that indicate an evolutionary process of streamlining similar to other dominant marine microbes such as members of the alphaproteobacterial *Pelagibacterales* (SAR11 clade) and the “*Ca*. Actinomarinales” (35, 38, 65). According to this theory which suggests that these modifications in the architecture of the genome represent a better evolutionary adaptation to oligotrophic environments, these microbes present a more cosmopolitan distribution compared to the other subclade. Despite their genomic divergence, the high similarity within the LGC group in the genomic features analyzed suggests that these genomes have reached the limit of their rationalization. Although the reduction in GC among other parameters does not lead to a reduction in genome size with respect to the other subclade, this genomic diversity provides an exceptional model for studying the evolutionary history of streamlined genomes. In the other subclade, there is a wide range of genomic architectures that may be due to different evolutionary histories or adaptations to different ecological niches. The presence of a genus (HGC2A) with similar characteristics to those of the LGC in terms of streamlining suggests that this evolutionary process may emerge in independent clades with parallel evolutionary trajectories. Although this study based on culture-independent approaches is a step further in understanding the population structure of this clade, genomic information obtained on the metabolic capabilities of these groups should be focused in future work on designing new isolation strategies to not only obtain more isolate strains but also to understand their role in aquatic environments. In addition, these findings provide a unique scenario to study the evolutionary processes related to genomic streamlining.

## MATERIAL AND METHODS

### Phylogenomic characterization

All available genomes belonging to the SAR116 clade were downloaded from the National Center for Biotechnology Information (NCBI), based on the Genome Taxonomy Database (GTDB) (33) (available up to August 2020) (table S1). CheckM (66) was used to estimate completeness and degree of contamination of the genomes and only those with completeness >50% and contamination <5% were kept. Phylophlan was used to establish the phylogenomic classification with a total of 258 genes shared among all suitable genomes (67). Along with the SAR116 genomes, a total of 85 reference genomes belonging to the SAR11 and Rickettsiales orders of the Alphaproteobacteria class were included. The resulting phylogenomic tree was analyzed and edited using iTOL (68).

### Genome comparison

For each genome, coding DNA sequences (CDS) of all genomes were predicted with Prodigal v2.6 (69). These sequences were annotated against the NCBI database of non-redundant protein sequences (NCBI’s NR) using DIAMOND (70) and against COG (71) and TIGFRAM (72) using HMMscan v3.1b2 (73). Subsequently, tRNA and rRNA genes were obtained using tRNAscan-SE v1.4 (74), ssu-align v0.1.1 (75) and meta-rna (76), respectively. To establish similarity and divergence of the genomes, the average nucleotide identity (ANI) between all the genomes were calculated using the JSpecies (77) package with standard parameters. Intrapopulation sequence diversity within each group was calculated using the average nucleotide identity calculate for metagenomics read with enveomics package (78) for R. In order to analyse streamlined genomic parameters GC content was calculated using the gecee program from the EMBOSS package (79). The number of paralogs was retrieved using cd-hit, iterating from 90 % to 30 %, in steps of 20% identity (80). Intergenic spacer size was calculated by measuring the distance between consecutive genes in all the genome. As a reference, we have included in the comparison representatives of well-known marine microbes: *Pelagibacter* sp. HTCC7211 (NCBI accession number GCA_000155895.1), *Candidatus* Actinomarina sp. AG-915-F11 (NCBI accession number GCA_902635395.1), *Alteromonas macleodii* ATCC27126 (NCBI accession number GCA_000172635.2), *Erythrobacter citreus* LAMA-915 (NCBI accession number GCA_001235865.1), *Synechococcus* sp. strain CC9902 (NCBI accession number GCA_000012505.1), “*Ca*. Nitrosopelagicus brevis” CN25 (NCBI accession number GCA_000812185.1), *Prochlorococcus marinus* MED4 (NCBI accession number GCA_000011465.1) and *Escherichia coli* str. a K-12 substr. MG1655 (NCBI accession number GCA_000005845.2).

In order to compare the genomic features of the genomes of the HGC and LGC subclades with several reference genomes, previously mentioned in this section, a principal component analysis (PCA) was performed using several genomic parameters: d*N*/d*S*, GC content, intergenic spacer and genome size as well as the number of paralogous genes. The FactoMineR (81) and factoextra (https://github.com/kassambara/factoextra) libraries of R were used for these analyses. The FactoMineR library was used to standardize the data during the PCA analysis. The plot was made using the Biplot function, in which values on the same side as the variable have a high value for that variable regardless of their position in the plot.

### Metagenomic fragment recruitment and SAR116 biogeography

Metagenomes from *Tara* Oceans expedition (16) were used to study ecological distribution patterns of SAR116 genomes. Only those genomes recruiting at least three reads per kilobase of genome and gigabase of metagenome (RPKG) and genome coverage of >70% and with an identity threshold of ≥ 98% were kept for further analyses. To avoid the bias caused by the high similarity rRNA operon, it was removed from all genomes before recruitment (35, 38). Metagenomic reads were aligned using BLASTN (82), using a cut-off of 98% nucleotide identity over a minimum alignment length of 50 nucleotides and ≥ 50% of each genome should be covered by reads for consideration. The same high-quality parameters were used for the metagenomic linear recruitment. The resulting alignments, together with the distribution of the reads according to the identity of the alignment (histogram) were plotted using the ggplot2 package in R.

### Functional classification

Since most of the genomes used are incomplete (MAGs and SAGs) we decided to use the pangenome of each of the established genera in order to compare them at the functional level. Pangenomes were generated using cd-hit (80) with a minimum percentage of identity of 70%, as well as coverage of at least 50%. The resulting pangenomes were annotated against three databases, SEED using DIAMOND (70) (40% identity and coverage greater than 50%), CAZy (83) using dbCAN (84) (HMMER Mode, e-value 10^−15^ and coverage greater than 35%) and KEGG (85) (KEGG Mapper, Reconstruct Brite, KEGG Orthology) using BlastKoala tool (86). We added in the comparison one of the two culture genomes (IMCC1322) as a reference.

## ACKNOWLEDGEMENTS

This work was supported by grants “VIREVO” CGL2016-76273-P [AEI/FEDER, EU] (cofounded with FEDER funds) from the Spanish Ministerio de Economía, Industria y Competitividad, “HIDRAS3” PROMETEU/2019/009 from Generalitat Valenciana and 5top100-program of the Ministry for Science and Education of Russia to FRV. JHM was supported by a Ph.D. fellowship from the Spanish Ministerio de Economía y Competitividad (BES-2014-067828)

## AUTHORS’ CONTRIBUTIONS

MLP and JHM conceived the study. JHM, JRG, LAH and MLP analysed the data. JHM, JRG, FRV and MLP contributed to write the manuscript.

## SUPPLEMENTARY MATERIAL

**Fig. S1**: Pairwise comparison among the SAR116 genomes using average nucleotide identity (ANI). Rectangles delimit subclades and genera, respectively.

**Fig. S2**. Linear recruitment plots of representative genomes of (A) genomospecies LCG1-A1 (B) and LCG2-C1 in two metagenomes. (C) An overview of the characteristic metabolism encoded in the flexible metagenomic island found in representatives within LCG2-C1.

**Fig. S3**. Genomic alignment (in amino acids) of the (A) different versions of the respiratory complex I (*nuo* and *nqr* operon) found in SAR116 genomes. *nqr* operon of Methylophilalles was added for the comparison. (B) Comparison of the proteorhodopsin gene cluster that includes genes involved in the retinal synthesis

**Table S1**: Detailed information about the genomes used in this study.

**Table S2**: Comparison of amino acid usage in SAR116 genera

**Table S3**. Genomic features of the SAR116 genomes genus versus reference genomes

## REFERENCES

1. Field CB, Behrenfeld MJ, Randerson JT, Falkowski P. 1998. Primary production of the biosphere: integrating terrestrial and oceanic components. Science 281:237–40.

2. Sogin ML, Morrison HG, Huber JA, Welch DM, Huse SM, Neal PR, Arrieta JM, Herndl GJ. 2006. Microbial Diversity in the Deep Sea and the Underexplored “Rare Biosphere,” p. 12115–12120. In Proceedings of the National Academy of Sciences. National Academy of Sciences.

3. Buchan A, LeCleir GR, Gulvik CA, González JM. 2014. Master recyclers: features and functions of bacteria associated with phytoplankton blooms. Nat Rev Microbiol. Nature Publishing Group.

4. Gómez-Pereira PR, Schüler M, Fuchs BM, Bennke C, Teeling H, Waldmann J, Richter M, Barbe V, Bataille E, Glöckner FO, Amann R. 2012. Genomic content of uncultured Bacteroidetes from contrasting oceanic provinces in the North Atlantic Ocean. Environ Microbiol 14:52–66.

5. Arandia-Gorostidi N, Weber PK, Alonso-Sáez L, Morán XAG, Mayali X. 2017. Elevated temperature increases carbon and nitrogen fluxes between phytoplankton and heterotrophic bacteria through physical attachment. ISME J 11:641–650.

6. López-Pérez M, Gonzaga A, Martin-Cuadrado A-BB, Onyshchenko O, Ghavidel A, Ghai R, Rodriguez-Valera F. 2012. Genomes of surface isolates of Alteromonas macleodii: The life of a widespread marine opportunistic copiotroph. Sci Rep 2:1–11.

7. Azam F, Malfatti F. 2007. Microbial structuring of marine ecosystems. Nat Rev Microbiol 5:782–791.

8. López-Pérez M, Jayakumar JM, Haro-Moreno JM, Zaragoza-Solas A, Reddi G, Rodriguez-Valera F, Shapiro OH, Alam M, Almagro-Moreno S. 2019. Evolutionary model of cluster divergence of the emergent marine pathogen vibrio vulnificus: From genotype to ecotype. MBio 10.

9. Wagner-Döbler I, Biebl H. 2006. Environmental Biology of the Marine Roseobacter Lineage. Annu Rev Microbiol 60:255–280.

10. Hou S, López-Pérez M, Pfreundt U, Belkin N, Stüber K, Huettel B, Reinhardt R, Berman-Frank I, Rodriguez-Valera F, Hess WR. 2018. Benefit from decline: the primary transcriptome of Alteromonas macleodii str. Te101 during Trichodesmium demise. ISME J https://doi.org/10.1038/s41396-017-0034-4.

11. Mullins TD, Britschgi TB, Krest RL, Giovannoni SJ. 1995. Genetic comparisons reveal the same unknown bacterial lineages in Atlantic and Pacific bacterioplankton communities. Limnol Oceanogr 40:148–158.

12. Giovannoni SJ, Britschgi TB, Moyer CL, Field KG. 1990. Genetic diversity in Sargasso Sea bacterioplankton. Nature 345:60–63.

13. Giovannoni SJ, Tripp HJ, Givan S, Podar M, Vergin KL, Baptista D, Bibbs L, Eads J, Richardson TH, Noordewier M, Rappé MS, Short JM, Carrington JC, Mathur EJ. 2005. Genome streamlining in a cosmopolitan oceanic bacterium. Science (80-) 309:1242–5.

14. Delong EF, Preston CM, Mincer T, Rich V, Hallam SJ, Frigaard N, Martinez A, Sullivan MB, Edwards R, Brito BR, Chisholm SW, Karl DM. 2006. Community Genomics among microbial assemblages in the Ocean’ s Interior. Science (80-) 311:496–503.

15. Rusch DB, Halpern AL, Sutton G, Heidelberg KB, Williamson S, Yooseph S, Wu D, Eisen JA, Hoffman JM, Remington K, Beeson K, Tran B, Smith H, Baden-Tillson H, Stewart C, Thorpe J, Freeman J, Andrews-Pfannkoch C, Venter JE, Li K, Kravitz S, Heidelberg JF, Utterback T, Rogers Y-H, Falcón LI, Souza V, Bonilla-Rosso G, Eguiarte LE, Karl DM, Sathyendranath S, Platt T, Bermingham E, Gallardo V, Tamayo-Castillo G, Ferrari MR, Strausberg RL, Nealson K, Friedman R, Frazier M, Venter JC. 2007. The Sorcerer II Global Ocean Sampling expedition: northwest Atlantic through eastern tropical Pacific. PLoS Biol 5:e77.

16. Sunagawa S, Coelho LP, Chaffron S, Kultima JR, Labadie K, Salazar G, Djahanschiri B, Zeller G, Mende DR, Alberti A, Cornejo-Castillo FM, Costea PI, Cruaud C, D’Ovidio F, Engelen S, Ferrera I, Gasol JM, Guidi L, Hildebrand F, Kokoszka F, Lepoivre C, Lima-Mendez G, Poulain J, Poulos BT, Royo-Llonch M, Sarmento H, Vieira-Silva S, Dimier C, Picheral M, Searson S, Kandels-Lewis S, Bowler C, de Vargas C, Gorsky G, Grimsley N, Hingamp P, Iudicone D, Jaillon O, Not F, Ogata H, Pesant S, Speich S, Stemmann L, Sullivan MB, Weissenbach J, Wincker P, Karsenti E, Raes J, Acinas SG, Bork P. 2015. Ocean plankton. Structure and function of the global ocean microbiome. Science 348:1261359.

17. Thorpe J, Stewart C, Venter JECEC, Smith H, Nealson K, Eisen JA, Platt T, Williamson S, Sutton G, Freeman J, Strausberg RL, Falcón LI, Bonilla-Rosso GG, Li K, Ferrari MR, Kravitz S, Gallardo V, Andrews-Pfannkoch C, Heidelberg JF, Wu D, Bermingham E, Frazier M, Yooseph S, Hoffman JM, Eguiarte LE, Friedman R, Baden-Tillson H, Venter JECEC, Utterback T, Heidelberg KB, Tran B, Beeson K, Karl DM, Sathyendranath S, Rogers Y-HH, Souza V, Rusch DB, Tamayo-Castillo G, Remington K, Halpern AL, Sutton G, Heidelberg KB, Williamson S, Yooseph S, Wu D, Eisen JA, Hoffman JM, Remington K, Beeson K, Tran B, Smith H, Baden-Tillson H, Stewart C, Thorpe J, Freeman J, Andrews-Pfannkoch C, Venter JECEC, Li K, Kravitz S, Heidelberg JF, Utterback T, Rogers Y-HH, Falcón LI, Souza V, Bonilla-Rosso GG, Eguiarte LE, Karl DM, Sathyendranath S, Platt T, Bermingham E, Gallardo V, Tamayo-Castillo G, Ferrari MR, Strausberg RL, Nealson K, Friedman R, Frazier M, Venter JECEC. 2007. The Sorcerer II Global Ocean Sampling expedition: Northwest Atlantic through eastern tropical Pacific. PLoS Biol 5:0398–0431.

18. Haro-Moreno JM, López-Pérez M, de la Torre JR, Picazo A, Camacho A, Rodriguez-Valera F. 2018. Fine metagenomic profile of the Mediterranean stratified and mixed water columns revealed by assembly and recruitment. Microbiome 6:128.

19. Pachiadaki MG, Brown JM, Brown J, Bezuidt O, Berube PM, Biller SJ, Poulton NJ, Burkart MD, La Clair JJ, Chisholm SW, Stepanauskas R. 2019. Charting the Complexity of the Marine Microbiome through Single-Cell Genomics. Cell 179:1623-1635.e11.

20. Rappé MS, Connon SA, Vergin KL, Giovannoni SJ. 2002. Cultivation of the ubiquitous SAR11 marine bacterioplankton clade. Nature 418:630–633.

21. Levin PA, Angert ER. 2015. Small but mighty: Cell size and bacteria. Cold Spring Harb Perspect Biol 7:1–11.

22. Kirchman DL. 2016. Growth Rates of Microbes in the Oceans. Ann Rev Mar Sci 8:285–309.

23. Giovannoni SJ, Cameron Thrash J, Temperton B. 2014. Implications of streamlining theory for microbial ecology. ISME J 8:1553–1565.

24. Dupont CL, Rusch DB, Yooseph S, Lombardo MJ, Alexander Richter R, Valas R, Novotny M, Yee-Greenbaum J, Selengut JD, Haft DH, Halpern AL, Lasken RS, Nealson K, Friedman R, Craig Venter J. 2012. Genomic insights to SAR86, an abundant and uncultivated marine bacterial lineage. ISME J 6:1186–1199.

25. Mizuno CM, Rodriguez-Valera F, Ghai R. 2015. Genomes of planktonic acidimicrobiales: Widening horizons for marine actinobacteria by metagenomics. MBio 6:e02083–14.

26. Giovannoni SJ, Rappé M. 2000. Evolution, diversity, and molecular ecology of marine prokaryotes. Microb Ecol Ocean John Wiley Sons, Inc, New York 47–84.

27. Giovannoni SJ, Vergin KL. 2012. Seasonality in Ocean Microbial Communities. Science (80-) 335:671–676.

28. Oh HM, Kwon KK, Kang I, Kang SG, Lee JH, Kim SJ, Cho JC. 2010. Complete genome sequence of “Candidatus puniceispirillum marinum” IMCC1322, a representative of the SAR116 clade in the Alphaproteobacteria. J Bacteriol 192:3240–3241.

29. Grote J, Bayindirli C, Bergauer K, Carpintero de Moraes P, Chen H, D’Ambrosio L, Edwards B, Fernández-Gómez B, Hamisi M, Logares R, Nguyen D, Rii YM, Saeck E, Schutte C, Widner B, Church MJ, Steward GF, Karl DM, DeLong EF, Eppley JM, Schuster SC, Kyrpides NC, Rappé MS. 2011. Draft genome sequence of strain HIMB100, a cultured representative of the SAR116 clade of marine Alphaproteobacteria. Stand Genomic Sci 5:269–278.

30. Choi DH, Park KT, An SM, Lee K, Cho JC, Lee JH, Kim D, Jeon D, Noh JH. 2015. Pyrosequencing revealed sar116 clade as dominant dddp-containing bacteria in oligotrophic nw pacific ocean. PLoS One 10.

31. Parks DH, Rinke C, Chuvochina M, Chaumeil P-A, Woodcroft BJ, Evans PN, Hugenholtz P, Tyson GW. 2017. Recovery of nearly 8,000 metagenome-assembled genomes substantially expands the tree of life. Nat Microbiol 903:1–10.

32. Bowers RM, Kyrpides NC, Stepanauskas R, Harmon-Smith M, Doud D, Reddy TBK, Schulz F, Jarett J, Rivers AR, Eloe-Fadrosh EA, Tringe SG, Ivanova NN, Copeland A, Clum A, Becraft ED, Malmstrom RR, Birren B, Podar M, Bork P, Weinstock GM, Garrity GM, Dodsworth JA, Yooseph S, Sutton G, Glöckner FO, Gilbert JA, Nelson WC, Hallam SJ, Jungbluth SP, Ettema TJG, Tighe S, Konstantinidis KT, Liu WT, Baker BJ, Rattei T, Eisen JA, Hedlund B, McMahon KD, Fierer N, Knight R, Finn R, Cochrane G, Karsch-Mizrachi I, Tyson GW, Rinke C, Lapidus A, Meyer F, Yilmaz P, Parks DH, Eren AM, Schriml L, Banfield JF, Hugenholtz P, Woyke T. 2017. Minimum information about a single amplified genome (MISAG) and a metagenome-assembled genome (MIMAG) of bacteria and archaea. Nat Biotechnol. Nature Publishing Group.

33. Parks DH, Chuvochina M, Waite DW, Rinke C, Skarshewski A, Chaumeil P-A, Hugenholtz P. 2018. A standardized bacterial taxonomy based on genome phylogeny substantially revises the tree of life. Nat Biotechnol 36:996.

34. López-Pérez M, Haro-Moreno JM, Coutinho FH, Martinez-Garcia M, Rodriguez-Valera F. 2020. The Evolutionary Success of the Marine Bacterium SAR11 Analyzed through a Metagenomic Perspective. mSystems 5:605–625.

35. López-Pérez M, Haro-Moreno JM, Iranzo J, Rodriguez-Valera F. 2020. Genomes of the “Candidatus Actinomarinales” Order: Highly Streamlined Marine Epipelagic Actinobacteria. mSystems 5:e01041–20.

36. Martinez-Gutierrez CA, Aylward FO. 2019. Strong Purifying Selection Is Associated with Genome Streamlining in Epipelagic Marinimicrobia. Genome Biol Evol 11:2887–2894.

37. Graham ED, Tully BJ. 2021. Marine Dadabacteria exhibit genome streamlining and phototrophy-driven niche partitioning. ISME J 15:1248–1256.

38. Haro-Moreno JM, Rodriguez-Valera F, Rosselli R, Martinez-Hernandez F, Roda-Garcia JJ, Gomez ML, Fornas O, Martinez-Garcia M, López-Pérez M. 2020. Ecogenomics of the SAR11 clade. Environ Microbiol 22:1748–1763.

39. Orellana LH, Ben Francis T, Krüger K, Teeling H, Müller MC, Fuchs BM, Konstantinidis KT, Amann RI. 2019. Niche differentiation among annually recurrent coastal Marine Group II Euryarchaeota. ISME J 13:3024–3036.

40. Rodriguez-Valera F, Martin-Cuadrado A-B, Rodriguez-Brito B, Pasić L, Thingstad TF, Rohwer F, Mira A. 2009. Explaining microbial population genomics through phage predation. Nat Rev Microbiol 7:828–36.

41. Reyes-Prieto A, Barquera B, Juárez O. 2014. Origin and Evolution of the Sodium -Pumping NADH: Ubiquinone Oxidoreductase. PLoS One 9:e96696.

42. Getz EW, Tithi SS, Zhang L, Aylward FO. 2018. Parallel Evolution of Genome Streamlining and Cellular Bioenergetics across the Marine Radiation of a Bacterial Phylum. MBio 9:e01089–18.

43. Kim AD, Baker AS, Dunaway-Mariano D, Metcalf WW, Wanner BL, Martin BM. 2002. The 2-Aminoethylphosphonate-Specific Transaminase of the 2-Aminoethylphosphonate Degradation Pathway. J Bacteriol 184:4134 LP – 4140.

44. Villarreal-Chiu J, Quinn J, McGrath J. 2012. The genes and enzymes of phosphonate metabolism by bacteria, and their distribution in the marine environment. Front Microbiol.

45. Sowell SM, Wilhelm LJ, Norbeck AD, Lipton MS, Nicora CD, Barofsky DF, Carlson CA, Smith RD, Giovanonni SJ. 2009. Transport functions dominate the SAR11 metaproteome at low-nutrient extremes in the Sargasso Sea. ISME J 3:93–105.

46. Olson DK, Yoshizawa S, Boeuf D, Iwasaki W, DeLong EF. 2018. Proteorhodopsin variability and distribution in the North Pacific Subtropical Gyre. ISME J https://doi.org/10.1038/s41396-018-0074-4.

47. Morris RM, Rappé MS, Connon SA, Vergin KL, Siebold WA, Carlson CA, Giovannoni SJ. 2002. SAR11 clade dominates ocean surface bacterioplankton communities. Nature 420:806–810.

48. Nakajima Y, Kojima K, Kashiyama Y, Doi S, Nakai R, Sudo Y, Kogure K, Yoshizawa S. 2020. Bacterium Lacking a Known Gene for Retinal Biosynthesis Constructs Functional Rhodopsins. Microbes Environ 35.

49. Sievert SM, Kiene RP, Schulz-Vogt HN. 2007. The sulfur cycle. Oceanography 20:117–123.

50. Crowe SA, Cox RP, Jones C, Fowle DA, Santibañez-Bustos JF, Ulloa O, Canfield DE. 2018. Decrypting the sulfur cycle in oceanic oxygen minimum zones. ISME J 12:2322–2329.

51. Glaubitz S, Kießlich K, Meeske C, Labrenz M, Jürgens K. 2013. SUP05 Dominates the gammaproteobacterial sulfur oxidizer assemblages in pelagic redoxclines of the central baltic and black seas. Appl Environ Microbiol 79:2767–2776.

52. Yoch DC. 2002. Dimethylsulfoniopropionate: Its Sources, Role in the Marine Food Web, and Biological Degradation to Dimethylsulfide. Appl Environ Microbiol 68:5804 LP – 5815.

53. Bullock HA, Luo H, Whitman WB. 2017. Evolution of Dimethylsulfoniopropionate Metabolism in Marine Phytoplankton and Bacteria. Front Microbiol.

54. González JM, Hernández L, Manzano I, Pedrós-Alió C. 2019. Functional annotation of orthologs in metagenomes: a case study of genes for the transformation of oceanic dimethylsulfoniopropionate. ISME J 13:1183–1197.

55. Kappler U, Schäfer H. 2014. Transformations of Dimethylsulfide BT - The Metal-Driven Biogeochemistry of Gaseous Compounds in the Environment, p. 279– 313. In Kroneck, PMH, Torres, MES (eds.),. Springer Netherlands, Dordrecht.

56. Reisch CR, Stoudemayer MJ, Varaljay VA, Amster IJ, Moran MA, Whitman WB. 2011. Novel pathway for assimilation of dimethylsulphoniopropionate widespread in marine bacteria. Nature 473:208–211.

57. Kang I, Oh H-M, Kang D, Cho J-C. 2013. Genome of a SAR116 bacteriophage shows the prevalence of this phage type in the oceans. Proc Natl Acad Sci 110:12343–12348.

58. Shah V, Zhao X, Lundeen RA, Ingalls AE, Nicastro D, Morris RM. 2019. Morphological Plasticity in a Sulfur-Oxidizing Marine Bacterium from the SUP05 Clade Enhances Dark Carbon Fixation. MBio 10:e00216–19.

59. Savoie ER, Lanclos VC, Henson MW, Cheng C, Getz EW, Barnes SJ, LaRowe DE, Rappé MS, Thrash JC. 2021. Ecophysiology of the cosmopolitan OM252 bacterioplankton (Gammaproteobacteria). bioRxiv 2021.03.09.434695.

60. Moran MA, Buchan A, González JM, Heidelberg JF, Whitman WB, Kiene RP, Henriksen JR, King GM, Belas R, Fuqua C, Brinkac L, Lewis M, Johri S, Weaver B, Pai G, Eisen JA, Rahe E, Sheldon WM, Ye W, Miller TR, Carlton J, Rasko DA, Paulsen IT, Ren Q, Daugherty SC, Deboy RT, Dodson RJ, Durkin AS, Madupu R, Nelson WC, Sullivan SA, Rosovitz MJ, Haft DH, Selengut J, Ward N. 2004. Genome sequence of Silicibacter pomeroyi reveals adaptations to the marine environment. Nature 432:910–913.

61. Poretsky RS, Hewson I, Sun S, Allen AE, Zehr JP, Moran MA. 2009. Comparative day/night metatranscriptomic analysis of microbial communities in the North Pacific subtropical gyre. Environ Microbiol 11:1358–1375.

62. Tuttle JH, Jannasch HW. 1977. Thiosulfate stimulation of microbial dark assimilation of carbon dioxide in shallow marine waters. Microb Ecol 4:9–25.

63. Ghosh W, Dam B. 2009. Biochemistry and molecular biology of lithotrophic sulfur oxidation by taxonomically and ecologically diverse bacteria and archaea. FEMS Microbiol Rev 33:999–1043.

64. van Vliet DM, von Meijenfeldt FAB, Dutilh BE, Villanueva L, Sinninghe Damsté JS, Stams AJM, Sánchez-Andrea I. 2020. The bacterial sulfur cycle in expanding dysoxic and euxinic marine waters. Environ Microbiol n/a.

65. Giovannoni SJ. 2017. SAR11 Bacteria: The Most Abundant Plankton in the Oceans. Ann Rev Mar Sci 9:231–255.

66. Parks DH, Imelfort M, Skennerton CT, Hugenholtz P, Tyson GW. 2015. CheckM: assessing the quality of microbial genomes recovered from isolates, single cells, and metagenomes. Genome Res 25:1043–55.

67. Segata N, Börnigen D, Morgan XC, Huttenhower C. 2013. PhyloPhlAn is a new method for improved phylogenetic and taxonomic placement of microbes. Nat Commun 4:2304.

68. Letunic I, Bork P. 2016. Interactive tree of life (iTOL) v3: an online tool for the display and annotation of phylogenetic and other trees. Nucleic Acids Res 44:W242–W245.

69. Hyatt D, Chen G-L, Locascio PF, Land ML, Larimer FW, Hauser LJ. 2010. Prodigal: prokaryotic gene recognition and translation initiation site identification. BMC Bioinformatics 11:119.

70. Buchfink B, Xie C, Huson DH. 2015. Fast and sensitive protein alignment using DIAMOND. Nat Methods 12:59–60.

71. Tatusov RL, Natale DA, Garkavtsev I V, Tatusova TA, Shankavaram UT, Rao BS, Kiryutin B, Galperin MY, Fedorova ND, Koonin E V. 2001. The COG database: new developments in phylogenetic classification of proteins from complete genomes. Nucleic Acids Res 29:22–8.

72. Haft DH, Loftus BJ, Richardson DL, Yang F, Eisen JA, Paulsen IT, White O. 2001. TIGRFAMs: a protein family resource for the functional identification of proteins. Nucleic Acids Res 29:41–43.

73. Eddy SR. 2011. Accelerated profile HMM searches. PLoS Comput Biol 7:e1002195.

74. Lowe TM, Eddy SR. 1996. TRNAscan-SE: A program for improved detection of transfer RNA genes in genomic sequence. Nucleic Acids Res 25:955–964.

75. Nawrocki EP. 2009. Structural RNA Homology Search and Alignment Using Covariance Models. Washington University.

76. Huang Y, Gilna P, Li W. 2009. Identification of ribosomal RNA genes in metagenomic fragments. Bioinformatics 25:1338–1340.

77. Richter M, Rossello-Mora R. 2009. Shifting the genomic gold standard for the prokaryotic species definition. Proc Natl Acad Sci 106:19126–19131.

78. Rodriguez-r LM, Konstantinidis KT. 2016. The enveomics collection : a toolbox for specialized analyses of microbial genomes and metagenomes. Peer J Prepr https://doi.org/10.7287/peerj.preprints.1900v1.

79. Rice P, Longden I, Bleasby A. 2000. EMBOSS: The European Molecular Biology Open Software Suite. Trends Genet 16:276–277.

80. Huang Y, Niu B, Gao Y, Fu L, Li W. 2010. CD-HIT Suite: A web server for clustering and comparing biological sequences. Bioinformatics 26:680–682.

81. Lê S, Josse J, Husson F. 2008. FactoMineR: An R Package for Multivariate Analysis. J Stat Softw 25:1–18.

82. Altschul SF, Madden TL, Schäffer AA, Zhang J, Zhang Z, Miller W, Lipman DJ. 1997. Gapped BLAST and PSI-BLAST: A new generation of protein database search programs. Nucleic Acids Res 25:3389–3402.

83. Lombard V, Golaconda Ramulu H, Drula E, Coutinho PM, Henrissat B. 2014. The carbohydrate-active enzymes database (CAZy) in 2013. Nucleic Acids Res 42.

84. Yin Y, Mao X, Yang J, Chen X, Mao F, Xu Y. 2012. DbCAN: A web resource for automated carbohydrate-active enzyme annotation. Nucleic Acids Res 40.

85. Kanehisa M, Sato Y, Kawashima M, Furumichi M, Tanabe M. 2016. KEGG as a reference resource for gene and protein annotation. Nucleic Acids Res 44:D457–D462.

86. Kanehisa M, Sato Y, Morishima K. 2016. BlastKOALA and GhostKOALA: KEGG Tools for Functional Characterization of Genome and Metagenome Sequences. J Mol Biol 428:726–731.

